# Emerging variants of concern in SARS-CoV-2 membrane protein: a highly conserved target with potential pathological and therapeutic implications

**DOI:** 10.1101/2021.03.11.434758

**Authors:** Lishuang Shen, Jennifer Dien Bard, Timothy J. Triche, Alexander R. Judkins, Jaclyn A. Biegel, Xiaowu Gai

## Abstract

Mutations in the SARS-CoV-2 Membrane (M) gene are relatively uncommon. The M gene encodes the most abundant viral structural protein, and is implicated in multiple viral functions, including initial attachment to the host cell via heparin sulfate proteoglycan, viral protein assembly in conjunction with the N and E genes, and enhanced glucose transport. We have identified a recent spike in the frequency of reported SARS-CoV-2 genomes carrying M gene mutations. This is associated with emergence of a new sub-B.1 clade defined by the previously unreported M:I82T mutation within TM3, the third of three membrane spanning helices implicated in glucose transport. The frequency of this mutation increased in the USA from 0.014% in October 2020 to 1.62% in February 2021, a 116-fold change. While constituting 0.7% of the isolates overall, M:I82T sub-B.1 lineage accounted for 14.4% of B.1 lineage isolates in February 2021, similar to the rapid initial increase previously seen with the B.1.1.7 and B.1.429 lineages, which quickly became the dominant lineages in Europe and California over a period of several months. A similar increase in incidence was also noted in another related mutation, V70L, also within the TM2 transmembrane helix. The rapid emergence of this sub-B.1 clade with recurrent I82T mutation suggests that this M gene mutation is more biologically fit, perhaps related to glucose uptake during viral replication, and should be included in ongoing genomic surveillance efforts and warrants further evaluation for potentially increased pathogenic and therapeutic implications.

## Introduction

Genomic surveillance is critical for identification of SARS-CoV-2 variants of concern (VOCs) (1). Children’s Hospital Los Angeles (CHLA) has routinely sequenced all viral isolates from over 2,900 pediatric and adult COVID-19 cases since March 2020. Using the CHLA COVID-19 Analysis Research Database (CARD), we have routinely performed local genomic epidemiology and genomic surveillance of viral sequences submitted to GISAID and NCBI GenBank (2, 3, 4). This allowed us to identify SARS-CoV-2 haplotypes and their localized transmission patterns that arose early and became dominant (5), specifically the D614G S spike protein mutation that was unidentified prior to April but which was identified in 99.3% of viral isolates from our pediatric COVID-19 patients by June of 2020 (6). We also identified the potential association of phylogenetic clade 20C with more severe pediatric disease (6). Here we report a new VOC with a signature mutation in the M protein gene, an otherwise overlooked but potentially significant site of increasing numbers of mutations, reminiscent of accumulating mutations in the Spike gene of previously reported VOCs, such as B.1.1.7 and B.1.351.

## Methods

### Ethics approval

Study design conducted at Children’s Hospital Los Angeles was approved by the Institutional Review Board under IRB CHLA-16-00429.

### SARS-CoV-2 whole genome sequencing

Whole genome sequencing of the 2900 samples previously confirmed at Children’s Hospital Los Angeles to be positive for SARS-CoV-2 by reverse transcription-polymerase chain reaction (RT-PCR) between March, 2020 to February 2021 was performed as previously described (6).

### SARS-CoV-2 sequence and variant analysis

Full-length SARS-CoV-2 sequences have been periodically downloaded from GISAID and NCBI GenBank. They are combined with sequences from CHLA patients, annotated, and curated using bioinformatics tools previously described (5).

### Phylogenetic analysis

Phylogenetic analysis was conducted using the NextStrain phylogenetic pipeline (version 3.0.1) (https://nextstrain.org/). Mafft (v7.4) was used in multiple sequence alignment (7), IQ-Tree (multicore version 2.1.1 COVID-edition) and TreeTime version 0.7.6 were used to infer and time-resolve evolutionary trees, and reconstruct ancestral sequences and mutations (8, 9).

### Protein structure prediction

M protein structural predictions were carried out using the Missense3D service hosted online by the Imperial College London (http://www.sbg.bio.ic.ac.uk/~missense3d/) (10).

## Results

We evaluated 143,609 USA SARS-CoV-2 viral genomes, including 2,900 from our own patients, and 622,033 global viral genomes, reported to GISAID and NCBI GenBank through late-February 2021. By measuring the ratios of the genomes carrying at least one missense mutation and genomes carrying at least one synonymous mutation in comparison to the reference genome (NC_045512) we identified distinctively different mutation profiles over time across SARS-CoV-2 genes (**Figure 1**). To evaluate these profiles, we performed exhaustive permutations of all possible changes at each base pair position of the SARS-CoV-2 gene to estimate missense mutations that could occur by chance. Using this approach, we estimated missense mutations should occur at least 2.7 times more frequently than synonymous mutations in the Envelope (E) gene, 3.1 times more frequently in the M gene, and as much as 3.8 times in ORF6 gene.

**Figure 1.**
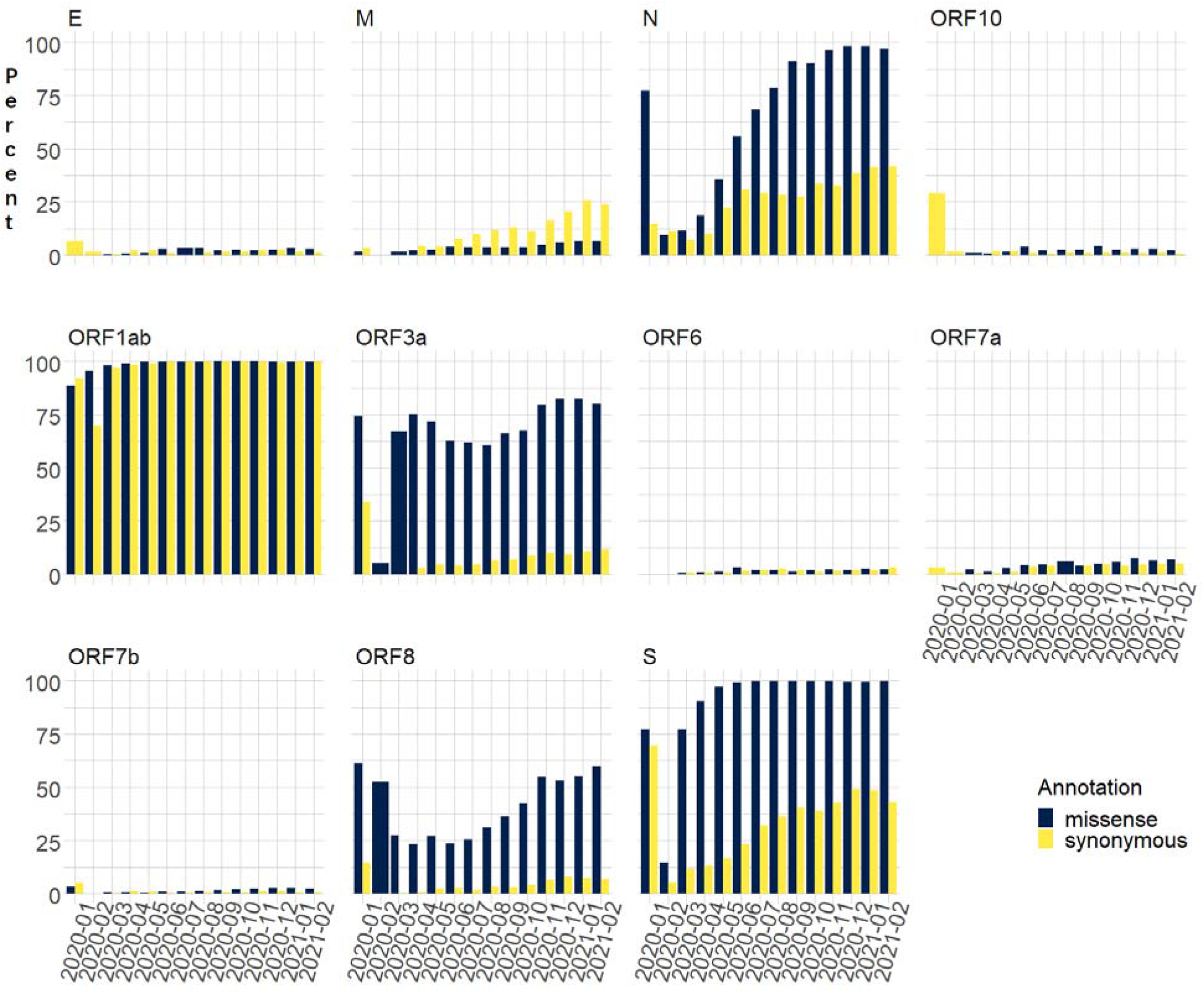
Percent of missense or synonymous mutations in viral isolates from USA over 14 months.

However, only the M gene has a ratio of missense to synonymous mutation carrying genomes consistently below 1.0 since the beginning of the pandemic. Further, this ratio has generally decreased over time. Among viral genomes from the USA as of late-February 2021, 6616 isolates (4.6%) carried missense and 22908 (16.0%) carried synonymous mutations in the M gene, for a ratio of 0.29. Globally, the ratio is even lower for the same period: there were 29,431 (4.73%) missense, and 197,205 (31.7%) synonymous mutations in the M gene, for a ratio of 0.149. This suggests that the M gene is highly conserved and potentially under strong purifying selection. It is thus of great potential interest that the incidence of some missense M gene mutations has recently increased over 100 fold in the past four months and continues to increase. The reason for this remains unclear but may suggest an underlying biologic advantage yet to be identified.

Other SARS-CoV-2 genes, notably the S (spike) gene and the large ORF1ab gene, appear to be tolerant of missense mutations, with multiple mutations in virtually every isolate. Indeed, some Spike missense mutations, like D614G and E484K, are advantageous, leading to rapid spread and increased frequency in the overall population and the emergence of a number of VOCs that uniformly include mutations like the D614G, which is now found in nearly every isolate worldwide but was unreported a year ago (11, 12). A different pattern is evident in ORF1ab where the relatively large size of this gene coupled with mutational tolerance has led to essentially all isolate showing one or more mutations in ORF1ab, producing a ratio close to 1.

For each M gene missense mutation, we calculated the percentage of mutation-carrying viral genomes at country and in USA state levels, and then compared the frequency increase vs previous months, to determine the timing and fold increase. The M gene is relatively silent, with only 4 missense mutations that together account for 0.4% or more of the global viral genomes in the month of February 2021 (M:A2S at 1.01%, M:V70L at 1.004%, M:I82T at 0.68%, and M:F28L at 0.41%). However, the percentage of viral genomes carrying missense M mutations has increased over time, and the accumulation of some mutations in the M protein appear to have surged recently both in the USA and globally. In the USA, 2.21%, 3.66% and 5.96% of reported viral genomes in April, August and December 2020 had missense M mutations. There has been a sharp increase in these missense mutations over the last three months, rising to 6.6% in the USA by February 2021 (**Figure 1**).

We identified six mutations which showed a significant increase in frequency, reaching 0.4% during the recent months and which could potentially account for this acceleration of M mutations (**Figure 2**). The M:I48V mutation is highly specific to the USA at 1.18% in January 2021 (**Table 1, Table S1**), 87 times that observed in samples from outside the USA (0.014%). Most (78.3%) of the mutation-carrying isolates belong to the B.1.375 lineage (**Table S3**). This mutation also shows considerable geographic variability, with the greatest frequency in isolates from the northeast and along the East coast: Rhode Island - 68.8%; Connecticut - 24.0%; New Hampshire - 17.8%; Florida - 15.9%; Massachusetts - 15.7%; Tennessee - 11.8%; Arkansas - 11.1% as of December 2020. Over the last three months, 129 of the 293 viral sequences from Rhode Island had the same M:I48V missense mutation, an approximately 9.5 fold higher frequency than in other USA locations. After peaking in December, the M:I48V missense mutation appear to be diminishing with a current 0.13% frequency in the USA (**Table 1, Table S1, Figure 2**).

**Figure 2.**
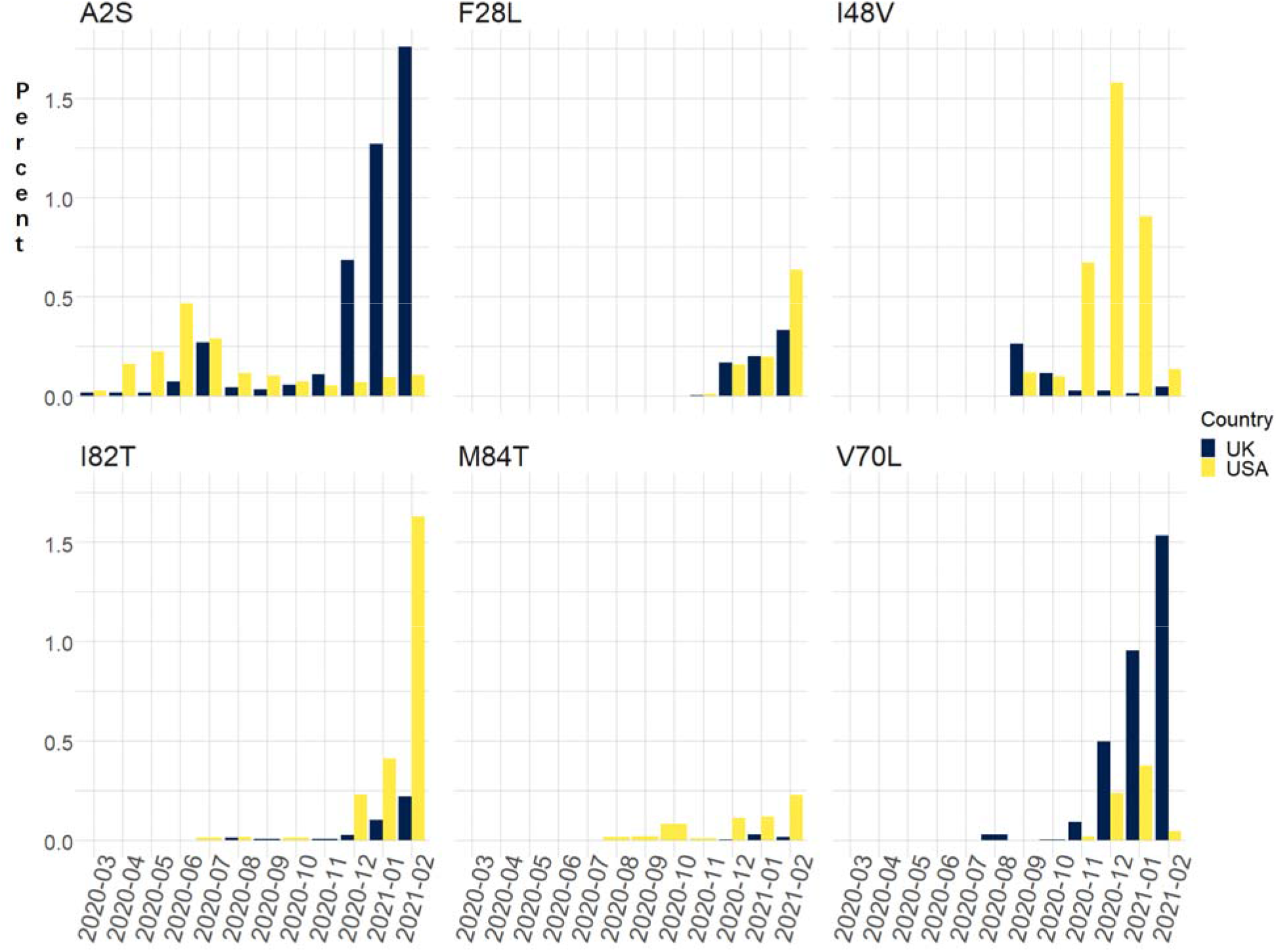
Percent of specific M mutations in viral isolates from USA and UK over 14 months.

In contrast, the M:I82T mutation increased in frequency 116 fold from 0.014% in October 2020 to 1.62% in February 2021 in the USA and continues to grow. While it predominately circulated in New York and New Jersey, over the past 2 months M:I82T has surged outside the USA including Aruba (5.2%) and Nigeria (33.1%) (**Table 1, Table S1**). This mutation presents mainly within the B.1 (44.0%) and B.1.525 (38.1%) lineages. Currently, 99.7% of the B.1.525 lineage isolates carry the M:I82T mutation. While this mutation is scattered across multiple phylogenetic clades, most cases cluster in two recent clades (**Figure 3, Figure 4**), suggesting a likely selective advantage in certain haplotype backgrounds.

**Figure 3.**
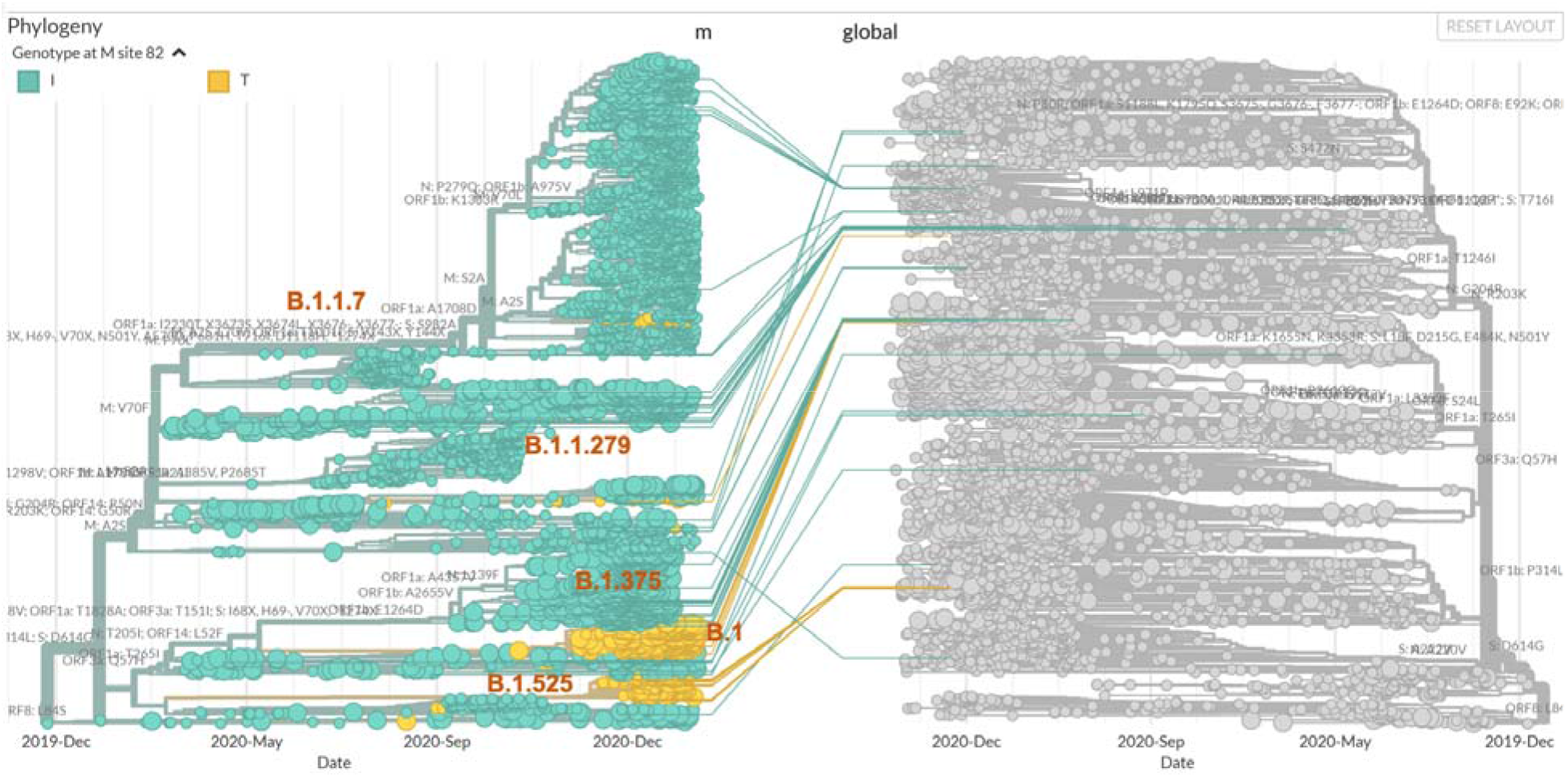
Phylogenetic tree of viral genomes carrying missense M mutations (left), colored by the genotypes at M:82 (I: green; T: yellow), overlaid on the global SARS-CoV-2 phylogenetic tree (right and grey).

**Figure 4.**
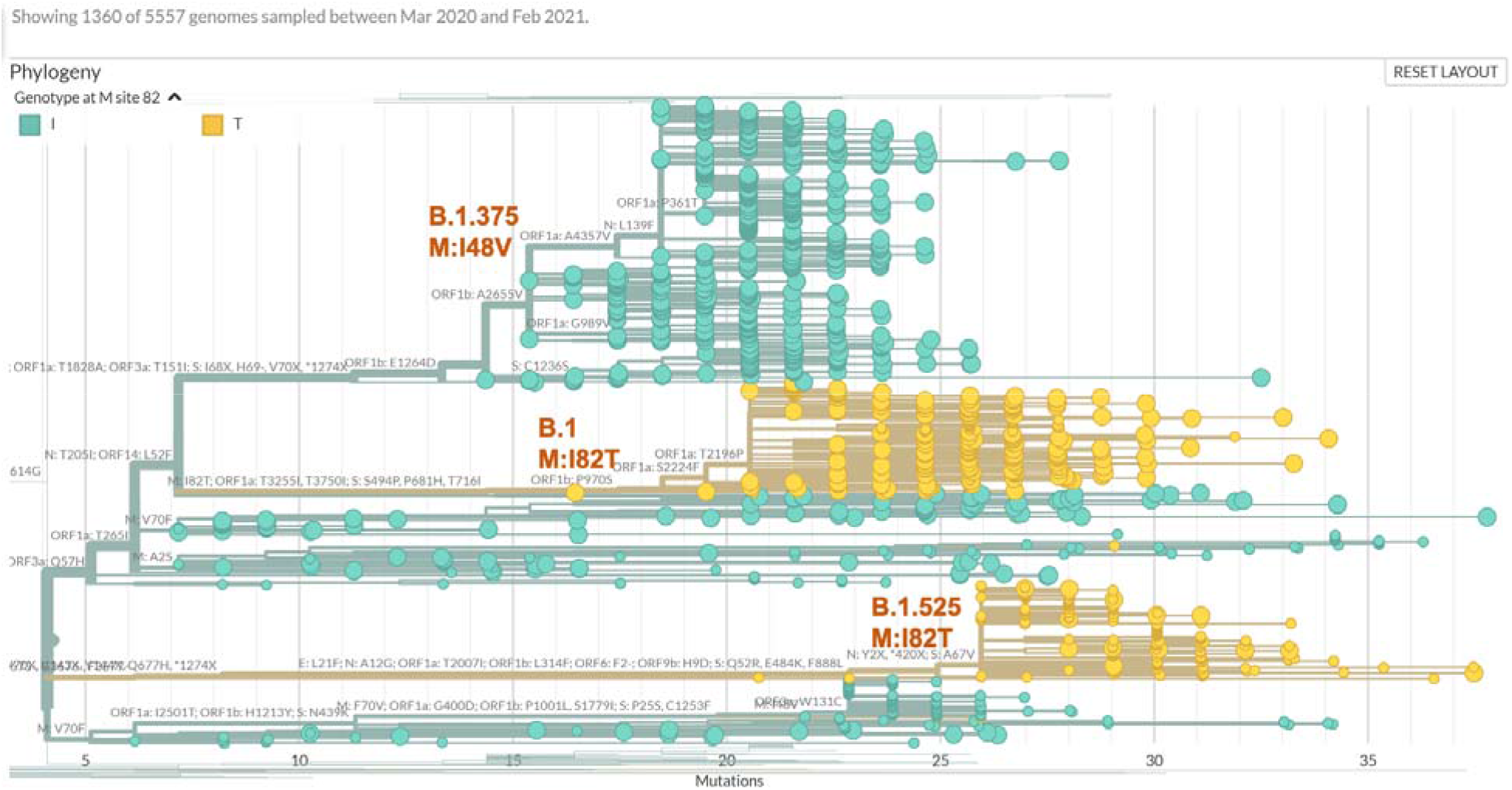
Detailed view of phylogenetic tree of viral genomes carrying missense M mutations colored by the genotypes at M:82 (I: green; T: yellow), demonstrating proposed new clade B.1 M:I82T (middle), falling between B.1.375 clade that carries M:I48V (top) and B.1.525 clade that also carries M:I82T (bottom).

The largest M:I82T carrying clade is part of a young M:I82T sub-B.1 lineage that has surged over the past 3 months to account for 14.4% of B.1 lineage isolates in February, and now constitutes 0.7% of all B.1 lineages. There were 10 other missense mutations present in at least 90% of the isolates in this clade and 8 of them were enriched by 73 to 146 fold compared to the general B.1 lineage including the 3 signature mutations in the spike protein (S:S494P, the S:P681H and S:T716I) found in the B.1.1.7 lineage (**Table 2)**. Thus, the M:I82T clade is significantly phylogenetically separated from other B.1 lineage clades, and may deserve consideration for a separate lineage designation (**Figure 3 & 4**).

This new sub-B.1 clade warrants close surveillance given the similarity to the initial patterns of B.1.1.7 and B.1.429, which quickly became the dominating lineages in Europe and California, respectively. The second largest M:I82T carrying clade arose only recently in December 2020 and is mainly circulating in Europe and Africa, where it was co-segregating with the S:E484K Spike protein mutation, forming lineage B.1.525 (**Table S2**, https://covlineages.org/global_report_B.1.525.html). This clade is smaller than the M:I82T clade in the USA, as described above, which lacks the S:E484K mutation. This suggests that M:I82T may confer a biologically selective advantage independent of the S:E484K, a known predictor of more severe viral infection. Another mutation carried by most isolates in this clade (98%) that is worth noting is N:T205I, because it is present in multiple VOCs including CAL.20C (B.1.429 and B.1.427) and B.1.351, and that M and N proteins are both important for viral assembly. The novel combination of M:I82T, the three signature Spike mutations (S:S494P, the S:P681H and S:T716I) from B.1.1.7, and the N:T205I mutation is therefore of particular concern.

Other M gene mutations have also increased in frequency. In the UK, M:V70L first appeared in September 2020 and the frequency increased 382 fold from 0.004% in October to 1.5% in February 2021, when it was also present in Switzerland at 3.6% and in Belgium at 3.0%. The M:V70L-carrying virus isolates are part of the minor lineages under the B.1.1.7 lineage. Another mutation in the same codon, M:V70F, has persisted at low frequencies across multiple countries since March 2020. M:F28L first appeared in November 2020 and is highest in Austria (02/2021, 39.3%), Ghana (01/2021, 6.4%) and Japan (01/2021, 2.3%) but also observed in Spain (2.1%), Belgium (0.4%), and the Netherlands (0.8%), by February 2021. Within the USA (0.6%), this mutation is present at high levels in isolates from Virginia (13.1%) and Maryland (3.8%). It presents mainly within the R. 1 (34.2%) and B.1.1.7 (48.9%) lineages, with 98% of the R. 1 lineage isolates carry the M:F28L as a signature mutation. M:A2S has existed widely across the world since last March, peaked to 0.9% in July globally, and has re-emerged globally at 1.0%, with levels of up to 3.22% in Spain and 1.76% in UK. Between November to February, about 87% (1537/1760) of the M:A2S -carrying virus isolates belong to the B.1.1.7 lineage. During the same period of October 2021 to February 2021, M:M84I increased from 0.1% to 0.23% in the USA. Further location and date details of these variants can be found in **Table S1**.

Focusing on the above-mentioned M mutations, we collected viral genomes from our own cases, GISAID, and GenBank that carried any of these mutations, as well as viral genomes that carried other M mutations by a haplotype similarity search which allowed a difference of up to five mutations across the genome than were the likely ancestral or descendant isolates in evolutionary context. This yielded 5,557 sequences that were analyzed for their phylogenetic relationships. The USA and UK sourced isolates were dominant in most clades, whereas limited mixtures exist in some cases (**Figure 5**). Cross checking the country of origin and Pangolin lineage assignment, we observed that many of the isolates belong to the B.1.1.7 lineage, while most I48V mutation isolates belonged to lineage B.1.375. The timing of recent increases in both the number and ratio of the M-gene mutation carrying isolates in Europe, especially UK, were most likely closely related to the B.1.1.7 (**Figure 6**). The B.1.1.7 lineage also carries a synonymous M mutation, and hence significantly reduced the ratio of missense to synonymous mutations in worldwide isolates. The M:V70L and M:A2S mutations both stayed within a narrow range of 1.1% to 1.5% within the B.1.1.7 lineage between December 2020 and February 2021, suggesting that a hijacking effect may account for these observed changes. However, the surge in other M mutations, and the emergence of a potentially new sub-B1 M:I82T carrying clade exceed what would be expected by a hijacking effect alone.

**Figure 5.**
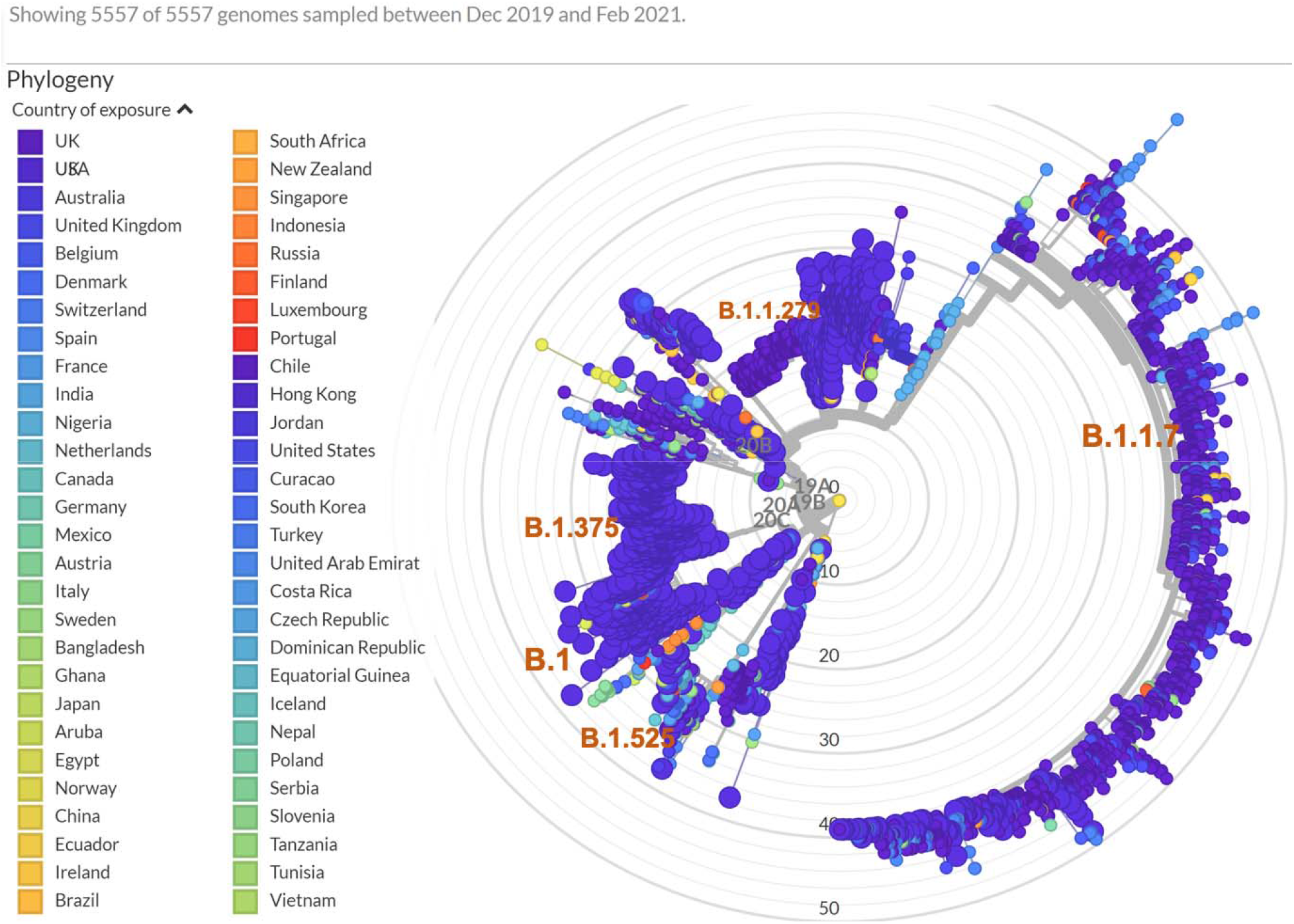
Phylogenetic analysis of viral genomes carrying missense M mutations, colored by country of origin. The USA viral genomes are represented by the larger dots to differentiate from the UK genomes.

**Figure 6.**
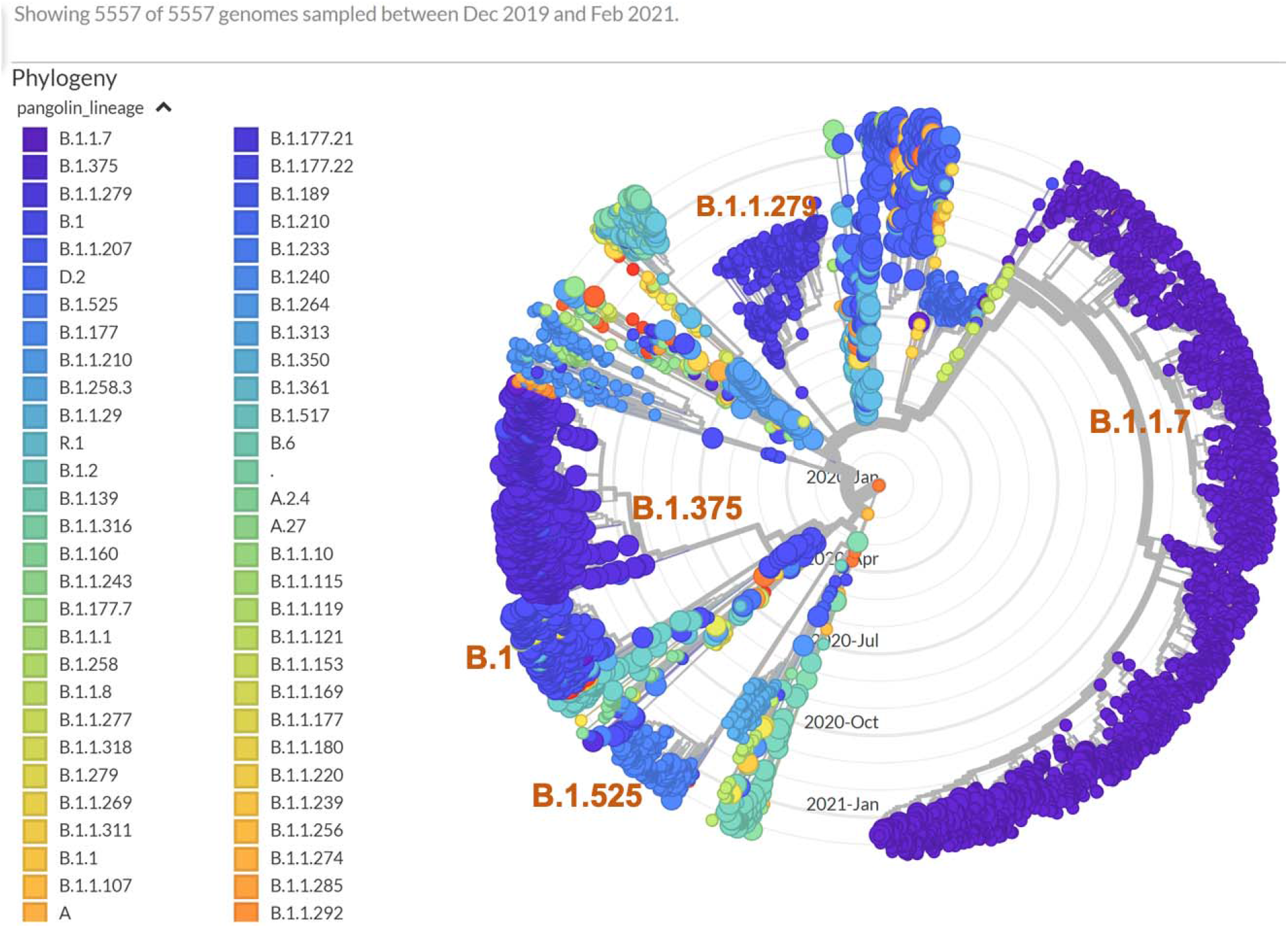
Phylogenetic analysis of viral genomes carrying missense M mutations, colored by lineage background.

Patient age information was collected for cases reported beginning in October and was divided into a group of cases carrying one of four M gene mutations (M:A2S, M:F28L, M:I82T, and M:V70L), which are the four more interesting mutations from above analysis. Frequencies of the other two mutations, M:I48V and M:M4T, were not increasing lately and hence excluded from this analysis. Each of the four M mutation groups had between 188-424 cases, and were compared against all the remaining cases without one of these M mutations, a total of 86,252 cases. One-way ANOVA analysis revealed significant difference in the patient age distribution among groups (p = 0.00092). Pairwise T-test, with Bonferroni multiple testing correction, indicates statistically significant patient age difference between each of the four M mutation carrying groups with the remaining group (adjusted p-values between 2.0e-4 to 9.5e-10), but not between each other (**Figure 7**). Mean patient ages of the four M-gene mutation groups were 4.6 - 6.3 years younger than the mean patient age in the “other” group (37.138.8 years compared to 43.4 years). We speculate that these M gene mutations may be associated with increased transmissibility among the younger population.

**Figure 7.**
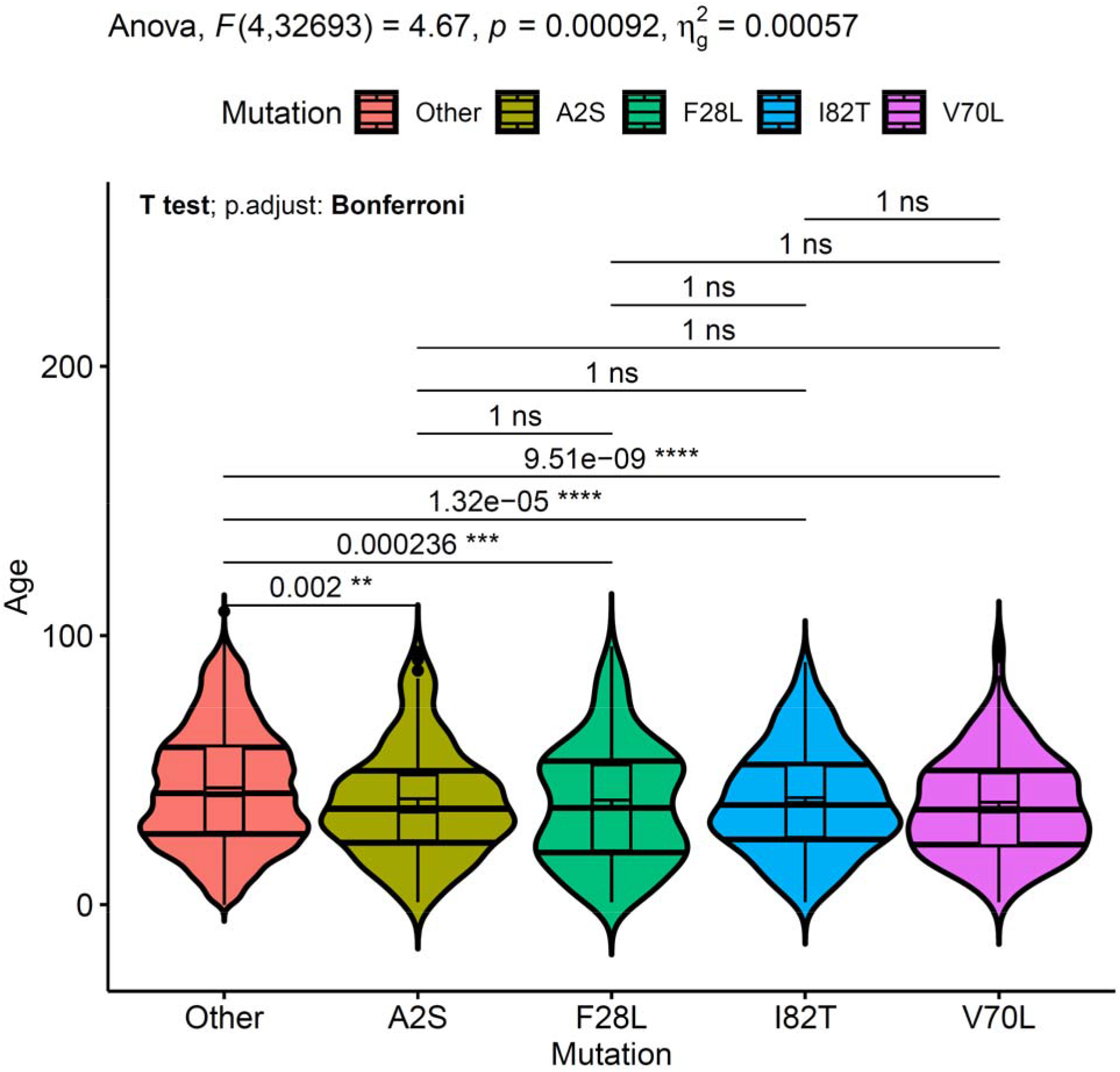
The ANOVA and t-test statistics between the five mutation groups. The plot was generated with R ggpubr package. Within each violin are the boxplot with error bars, and the horizontal lines at the quantiles 0.25, 0.50 and 0.75 of the density estimates. The pairwise t-test adjusted p-values and significance are shown to the upper part.

## Discussion

The M gene encodes the most abundant of three SARS-CoV-2 structural proteins, in this case a 222 amino acid protein that is highly conserved between SARS-CoV and SARS-CoV-2 (identity: 90.5%; similarity: 98.2%) (13). Comparatively little attention has been paid to the M protein in the COVID-19 pandemic literature but it is known to be important for viral assembly, and in addition it markedly inhibits type I and III interferon production and thus dramatically inhibits the innate immune response (14, 15). That in turn blunts the T-cell mediated immune response which is known to be important in overall immunity to SARS. In SARS due to SARS-CoV, the M protein is the dominant immunogen for T-cell response (16). In COVID-19 due to SARS-CoV-2, T-cell response has been identified as a critical determinant of outcome, with poor T-cell response to M protein epitopes found in patients with fatal outcome (17, 18). T-cell responses are a critical part of the successful immune response against emerging VOCs that may enable immune evasion (19, 20). Therapeutic strategies that target the M protein and thus modulate T-cell responses have recently been proposed as promising alternatives to current ones (21).

*In silico* analysis revealed that the M protein structure was similar to that of the glucose transporter SemiSWEET with three transmembrane helical domains, based upon which the M protein is thought to be involved in enhanced glucose transport in host cells with replicating virus, and thus may aid in rapid viral proliferation, replication, and immune evasion (22). The M:I82T mutation falls in the third transmembrane helical domain (22). These transmembrane domains vary in number in the SWEET - 7, SemiSWEET – 3, and GLUT1 - 14 glucose transport family and are thought to bind and transport glucose, yet another function of the M protein that also initiates viral binding to the cell membrane heparin sulfate proteoglycan (via the N terminal exposed fragment), viral protein assembly via the internal carboxyterminal fragment, and immune evasion by inhibition of nuclear transport of NFκB signal transducers involved in interferon induction. Structural prediction however, did not suggest significant impact of structural changes to be caused by the M:I82T mutation which is a hydrophobic to slightly polar amino acid change (**Figure 8**).

**Figure 8.**
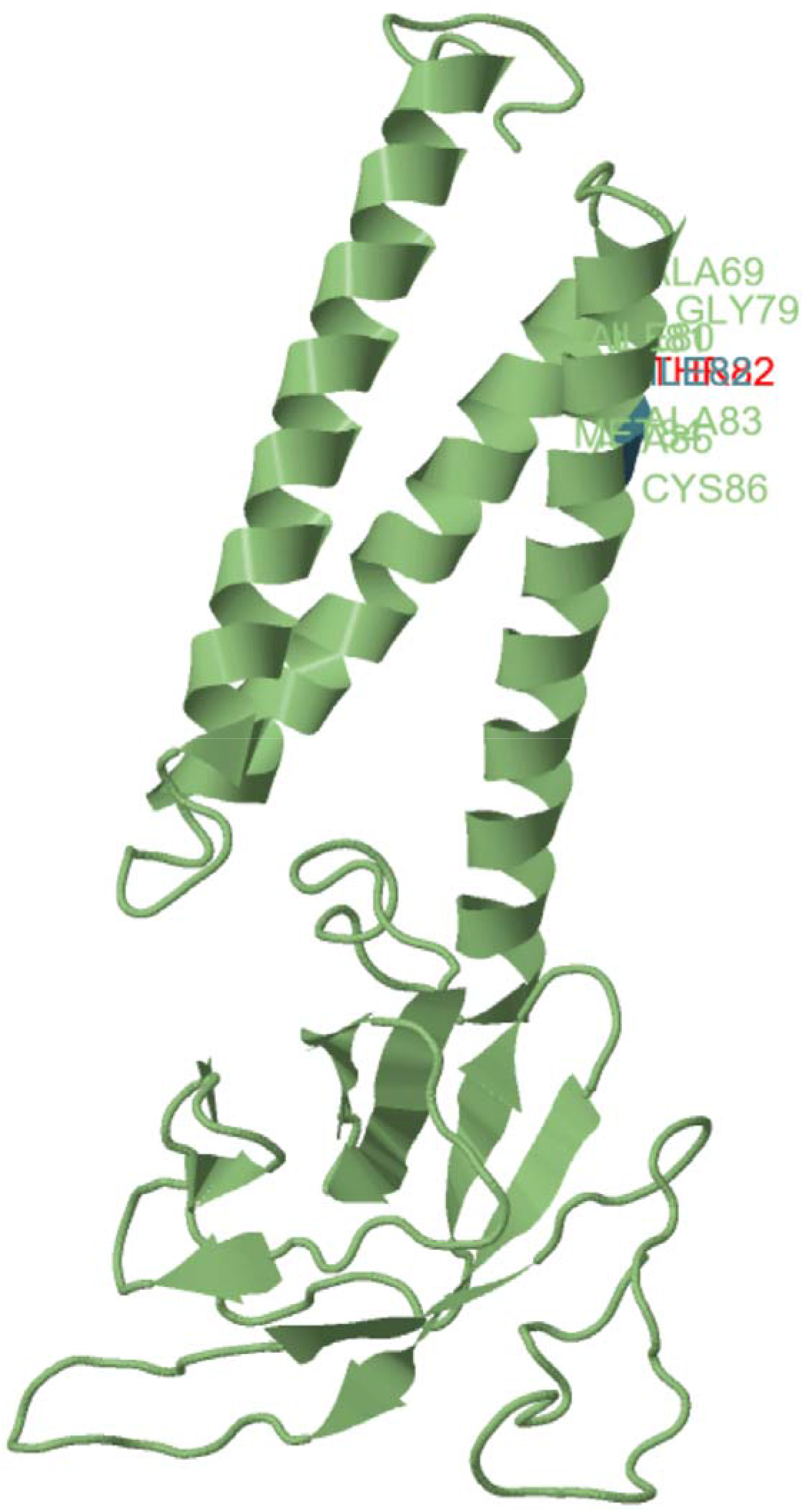
Missense3D prediction of 3D structures of the wide type (ILE82, green) and mutant (THR82, red) M proteins. (Based on PDB: qhd43419; https://zhanglab.ccmb.med.umich.edu/COVID-19/)

We have identified that the M gene, though otherwise highly conserved throughout most of the pandemic, is now undergoing rapidly increasing mutation with a recent surge in isolates carrying previously unreported M gene mutations in the USA and globally. In particular, we identified the emergence and rapid growth over the last three months of a novel M:I82T clade in the eastern USA and a V70L mutation in the UK. Both mutations involve the putative glucose transport transmembrane helices of the M protein. Isolates carrying the M:182T in the USA have increased from 0.014% in October 2020 to 1.62% in February 2021, a 116-fold change, and now represents 3.33% of isolates reported (February 16^th^ thru March 4^th^ 2021). We also noted a reduced average age for patients carrying these M missense mutations. Given its rapid emergence, and the role of the M protein in multiple critical viral functions including host cell binding, innate immune and T cell responses, immune evasion, and potential alterations in glucose transport, this novel M:I82T clade warrants inclusion in ongoing SARS-CoV-2 genomic surveillance and further evaluation for potential pathogenicity.

## Acknowledgments

We would like to acknowledge all members of the Department of Pathology and Laboratory Medicine (PLM) at Children’s Hospital Los Angeles for dedication towards providing excellent patient care throughout the pandemic, including especially the rapid launch of both the COVID-19 diagnostic test in the Clinical Microbiology and Virology Laboratory and the SARS-CoV-2 whole genome sequencing assay at the Center for Personalized Medicine in March 2020, both with the support of Thermo Fisher and Paragon Genomics, with whom these assays were developed. We would like to acknowledge the frontline healthcare workers who remain devoted in the fight against COVID-19. We would also like to acknowledge all institutions that have contributed the SARS-CoV-2 sequences timely, as well as NCBI, GISAID, and Nextstrain for providing valuable resources for SARS-CoV-2 genomics.

---

**Supplemental Table 1.** Summary of the frequencies of M gene missense mutations in isolates over 14 months.

| **Mutation** | **Month** | **# in USA** | **% in USA** | **% non-USA** | **% world** |
| --- | --- | --- | --- | --- | --- |
| M:V70L | 2021-02 | 5 | 0.047 | 1.255 | 1.004 |
| M:V70L | 2021-01 | 137 | 0.373 | 0.745 | 0.642 |
| M:V70L | 2020-12 | 41 | 0.237 | 0.302 | 0.288 |
| M:V70L | 2020-11 | 2 | 0.02 | 0.064 | 0.057 |
| M:F28L | 2021-02 | 67 | 0.631 | 0.355 | 0.413 |
| M:F28L | 2021-01 | 72 | 0.196 | 0.186 | 0.189 |
| M:F28L | 2020-12 | 27 | 0.156 | 0.115 | 0.124 |
| M:F28L | 2020-11 | 1 | 0.01 | 0.004 | 0.005 |
| M:I82T | 2021-02 | 172 | 1.62 | 0.435 | 0.683 |
| M:I82T | 2021-01 | 150 | 0.408 | 0.223 | 0.275 |
| M:I82T | 2020-12 | 40 | 0.231 | 0.043 | 0.082 |
| M:I82T | 2020-10 | 1 | 0.014 |  | 0.002 |
| M:I82T | 2020-08 | 1 | 0.016 | 0.004 | 0.007 |
| M:I82T | 2020-07 | 1 | 0.012 | 0.006 | 0.008 |
| M:I48V | 2021-02 | 14 | 0.132 | 0.037 | 0.057 |
| M:I48V | 2021-01 | 331 | 0.901 | 0.008 | 0.255 |
| M:I48V | 2020-12 | 275 | 1.592 | 0.02 | 0.345 |
| M:I48V | 2020-11 | 67 | 0.669 | 0.016 | 0.118 |
| M:I48V | 2020-10 | 7 | 0.097 | 0.075 | 0.078 |
| M:I48V | 2020-09 | 6 | 0.119 | 0.153 | 0.148 |
| M:I48V | 2020-01 | 1 | 1.613 |  | 0.148 |
| M:M84T | 2021-02 | 24 | 0.226 | 0.032 | 0.073 |
| M:M84T | 2021-01 | 43 | 0.117 | 0.025 | 0.05 |
| M:M84T | 2020-12 | 19 | 0.11 | 0.017 | 0.036 |
| M:M84T | 2020-11 | 1 | 0.01 | 0.009 | 0.009 |
| M:M84T | 2020-10 | 6 | 0.083 |  | 0.011 |
| M:M84T | 2020-09 | 1 | 0.02 | 0.004 | 0.006 |
| M:M84T | 2020-08 | 1 | 0.016 | 0.086 | 0.072 |
| A2S | 2021-02 | 11 | 0.104 | 1.248 | 1.01 |
| A2S | 2021-01 | 35 | 0.095 | 0.871 | 0.657 |
| A2S | 2020-12 | 12 | 0.069 | 0.467 | 0.384 |
| A2S | 2020-11 | 5 | 0.05 | 0.093 | 0.087 |
| A2S | 2020-10 | 5 | 0.07 | 0.055 | 0.057 |
| A2S | 2020-09 | 5 | 0.1 | 0.049 | 0.057 |
| A2S | 2020-08 | 7 | 0.115 | 0.289 | 0.253 |
| A2S | 2020-07 | 24 | 0.289 | 1.174 | 0.888 |
| A2S | 2020-06 | 52 | 0.467 | 0.349 | 0.401 |
| A2S | 2020-05 | 18 | 0.223 | 0.007 | 0.082 |
| A2S | 2020-04 | 19 | 0.158 | 0.039 | 0.069 |
| A2S | 2020-03 | 3 | 0.023 | 0.008 | 0.012 |

**Supplemental Table 2.** Potential signature mutations in the B.1 sub-clade that carries the M:I82T mutation

| # Isolates in clade | Mutations | Gene | Amino Acid Change | Annotation | # Isolates w/ mutation | % Isolates in clade w/ mutation |
| --- | --- | --- | --- | --- | --- | --- |
| 348 | **T26767C** | M | I82T | missense | 348 | 100 |
| 348 | **C28887T** | N | T205I | missense | 341 | 97.99 |
| 348 | **T23042C** | S | S494P | missense | 332 | 95.4 |
| 348 | **C23709T** | S | T716I | missense | 316 | 90.8 |
| 348 | **C23604A** | S | P681H | missense | 336 | 96.55 |
| 348 | **A23403G** | S | D614G | missense | 347 | 99.71 |
| 348 | **G25563T** | ORF3a | Q57H | missense | 332 | 95.4 |
| 348 | **A6851C** | orf1ab | T2196P | missense | 174 | 50 |
| 348 | **C16375T** | orf1ab | P5371S | missense | 328 | 94.25 |
| 348 | **C6936T** | orf1ab | S2224F | missense | 215 | 61.78 |
| 348 | **C11514T** | orf1ab | T3750I | missense | 327 | 93.97 |
| 348 | **C10029T** | orf1ab | T3255I | missense | 328 | 94.25 |
| 348 | **C1059T** | orf1ab | T265I | missense | 336 | 96.55 |
| 348 | **C14408T** | orf1ab | P4715L | missense | 343 | 98.56 |
| 348 | **A1180G** | orf1ab | P305P | synonymous | 317 | 91.09 |
| 348 | **T20748C** | orf1ab | Y6828Y | synonymous | 329 | 94.54 |
| 348 | **A1180G** | orf1ab | P305P | synonymous | 317 | 91.09 |
| 348 | **T20748C** | orf1ab | Y6828Y | synonymous | 329 | 94.54 |
| 348 | **C6730T** | orf1ab | N2155N | synonymous | 260 | 74.71 |
| 348 | **C3037T** | orf1ab | F924F | synonymous | 345 | 99.14 |
| 348 | **C241T** | 5UTR_orf1ab | . | upstream_gene | 343 | 98.56 |
| 348 | **C29719T** | ORF10_3UTR | . | intergenic | 318 | 91.38 |

## Notes

### Competing Interest Statement

The authors have declared no competing interest.

